# Immune priming in the insect gut: a dynamic response revealed by ultrastructural and transcriptomic changes

**DOI:** 10.1101/2024.11.21.624623

**Authors:** Moritz Baur, Nora K. E. Schulz, Lilo Greune, Zoe M. Länger, Jürgen Eirich, Iris Finkemeier, Robert Peuß, Petra Dersch, Joachim Kurtz

## Abstract

Research on forms of memory in innate immune systems has recently gained momentum with the study of trained immunity in vertebrates and immune priming in invertebrates. Immune priming provides protection against previously encountered pathogens. However, causes and mechanisms of immune priming are still not well understood in most organisms. In this work, we combine RNA sequencing with transmission electron microscopy to investigate the dynamic processes during priming in the gut of a well-established model for oral immune priming, consisting of the host *Tribolium castaneum* and its entomopathogen *Bacillus thuringiensis tenebrionis (Btt)*. We show that priming with specific, pathogen-derived virulence-relevant factors causes damage in the gut of *T. castaneum* larvae, which leads to an early physiological stress response as well as the upregulation of a specific set of immune genes. This response diminishes over time yet enables the gut to upregulate genes known to interfere with *Btt* virulence when *T. castaneum* larvae are later exposed to infectious *Btt* spores. The identification of these processes increases our understanding of immune priming as a dynamic process where cellular responses in concert with specific gene regulation prepare the gut tissue and thereby enable more efficient protection against infection. Such work can further help us understand the origin and mechanism of innate immune memory.

**Author summary:** Invertebrate animals do not possess a classical adaptive immune system. And yet, many of these animals show forms of immune memory, collectively called immune priming. In this work, we investigate the causes of oral immune priming in the red flour beetle *Tribolium castaneum* in response to its bacterial entomopathogen *Bacillus thuringiensis tenebrionis*. Primary exposure to *Btt* spent culture media supernatants enables the larvae of *T. castaneum* to better withstand a subsequent infection with infectious *Btt* spores. We show that exposing *T. castaneum* larvae to those supernatants leads to damage in the gut resulting in a strong stress and immune response early after priming and a targeted up-regulation of beneficial genes upon the secondary re-encounter with *Btt*. These results suggest fundamental stress responses can be involved in innate immune memory phenomena, which has implications in translational research across species.

## Introduction

Immune systems are a universal feature across the tree of life. As checks and balance systems that discriminate between self and non-self, they allow for protection against harmful pathogens and parasites while reducing self-harm. Immune systems of vertebrate animals have classically been divided into the innate and the adaptive arm. Invertebrates lack the specialized cellular machinery that is needed to mount an adaptive immune response in the classical sense [1, 2]. Over the past decades, this clear separation has been challenged by work on both vertebrate and invertebrate immunity. In vertebrates, innate immune memory that has been denoted ‘trained immunity’ acts through the rewiring of metabolism and epigenetic imprinting of immune and non-immune (e.g., epithelial) cells [3–6].

In invertebrates, numerous studies have described enhanced survival upon infection after a previous encounter with pathogens or their cues, despite the absence of the adaptive immune cellular machinery [7–12]. These observations, usually denoted as immune priming, are widespread across invertebrates and entail many distinct phenomena [13]. Systemic or septic immune priming is achieved by pricking or injecting a host with sublethal doses, inactivated microbes, or immunoreactive cues derived from those microbes before infecting it, in the same way, with living microbes. This route of priming can be specific, providing a survival benefit only if the priming agent is the same as the infecting agent [8, 12, 14–16], or unspecific, providing a benefit during a subsequent infection even if the infecting agent is different from the one used for priming [10]. Immune priming is also possible via the oral route. Like septic immune priming, oral immune priming can be specific depending on the host and microorganism involved [7, 17, 18]. The two different routes of immune priming (septic and oral) likely function via different mechanisms [19], although it has also been reported in some organisms that oral immune priming can protect a host against a subsequent septic infection [20, 21]. Immune priming further is transmittable to subsequent generations, a phenomenon termed transgenerational immune priming [22–25]. While mechanisms of septic immune priming have been studied over the past years, similar work on oral immune priming is still scarce.

Immune priming via the oral route is likely a complex process. The microbiome seems to play a role in some species [15, 26, 27]. In mosquitoes and bean bugs, members of the microbiome take an active part by breaching the gut epithelium, thereby stimulating immune responses that provide systemic immunization [15, 27]. In an important model for oral immune priming, the red flour beetle *Tribolium castaneum*, a shift in the microbiome composition upon priming has been described [28]. Oral immune priming enables insects to better resist a subsequent infection by ingested pathogens in an environment with high pathogen exposure. When the priming agent is taken up via the oral route, the intestine is the first tissue reacting to the presented stimuli.

In the present work, we focus on gut-associated processes of oral immune priming in *T. castaneum* when confronted with its natural entomopathogen *Bacillus thuringiensis tenebrionis* (*Btt*). *B. thuringiensis* is a spore-forming gram-positive bacterium. During sporulation, it produces crystal toxins (Cry toxins), which become solubilized upon ingestion by the host and form pores in the gut epithelium [29, 30]. Upon germination, *B. thuringiensis* produces a large number of enzymes and toxins to further damage its hosts’ gut tissue, ultimately leading to the death of the host [29, 30]. Once *B. thuringiensis* has exhausted the hosts’ nutrients, vegetative cells sporulate, ready to infect the next host [29, 30]. Greenwood et al. [31] found that oral immune priming of *T. castaneum* larvae with filter sterilized medium derived from sporulating *Btt* cells leads to a shift in whole-body gene expression profiles, which is still detectable four days later. Several immune genes remain up-regulated in primed hosts upon pathogen exposure, and some genes are distinctly induced by challenge in those animals that only received a prior priming [31]. Moreover, recent work comparing proteomes of filtered growth media supernatants derived from closely related *B. thuringiensis* isolates, that either do or do not lead to priming of *T. castaneum* larvae, enabled the identification of candidate virulence factors that might induce oral immune priming, including the plasmid-encoded Cry3Aa toxin [32].

We currently lack a comprehensive view of the associated ultrastructural and physiological changes in the gut that lead to the observed enhanced survival upon priming. Here, we studied ultrastructural changes using transmission electron microscopy together with changes in gene expression using RNA-seq of gut tissue of *T. castaneum* larvae at different time points after oral priming and infection to gain information about the mechanisms contributing to oral immune priming in *T. castaneum*. For these experiments, we made use of an improved control *Btt* strain for priming, which was cured of the Cry-toxin-carrying plasmid to identify bacterial candidate proteins leading to priming. As a result, we provide important insights into mechanisms that might constitute innate immune memory in invertebrates, in interplay with their natural pathogens.

## Materials and methods

### Model organisms

*T. castaneum* larvae used in this study stem from the Croatia 1 (Cro1) strain [28]. Cro1 was collected in 2010 in Croatia and maintained in large groups of non-overlapping generations since then. Larvae were reared on heat-sterilized (75°C for 24 h) wheat flour (Bio Weizenmehl Type 550, dm-drogerie markt GmbH + Co. KG) including 5% Brewer’s yeast at 30°C, 70% humidity and a 12 h day/night rhythm.

*Bacillus thuringiensis* bv *tenebrionis* (*Btt,* BGSCID4AA1) spores were purchased from the Bacillus Genetic Stock Center of the Ohio State University (USA). *Btt Δ p188* (previously denoted Btt – [32]), lacking the 188 kb plasmid, was obtained by serial passage of *Btt* at high temperatures [32].

### Study design

To understand the immediate responses following an oral priming treatment and which factors are involved in mounting a successful immune priming response, we combined proteomics data from priming media and larval size measurements, with electron microscopy and RNA-seq approaches (Fig 1). 14-day old *T. castaneum* larvae were individually exposed to either medium-(*Bt* growth medium), priming (*Btt*)- or control-(*Btt Δ p188*) flour mixes (Priming treatment) in a well of a 96-well plate. After 24 hours, larvae were transferred onto PBS flour mixes and left for another four days. Finally, larvae were transferred onto PBS- or *Btt* spore-flour mixes (Challenge treatment). For the size measurement (Fig 1), larvae were randomly sampled before exposure to the treatment diets, and the same larvae were re-sampled before exposure to the challenge treatment. For electron microscopy and RNA-sequencing (Fig 1), gut samples were taken from randomly sampled larvae at three hours and 24 hours post-exposure to the priming treatment as well as four hours post-exposure to the challenge treatment five days after the initial priming. In all experiments, the larvae were kept on the challenge treatment, and survival was recorded for three to five days post-exposure. Finally, the proteomes of spent culture media supernatants derived from *Btt* and *Btt Δ p188* cultures were compared to narrow down the agents causing the immune priming response [32].

**Fig 1:**
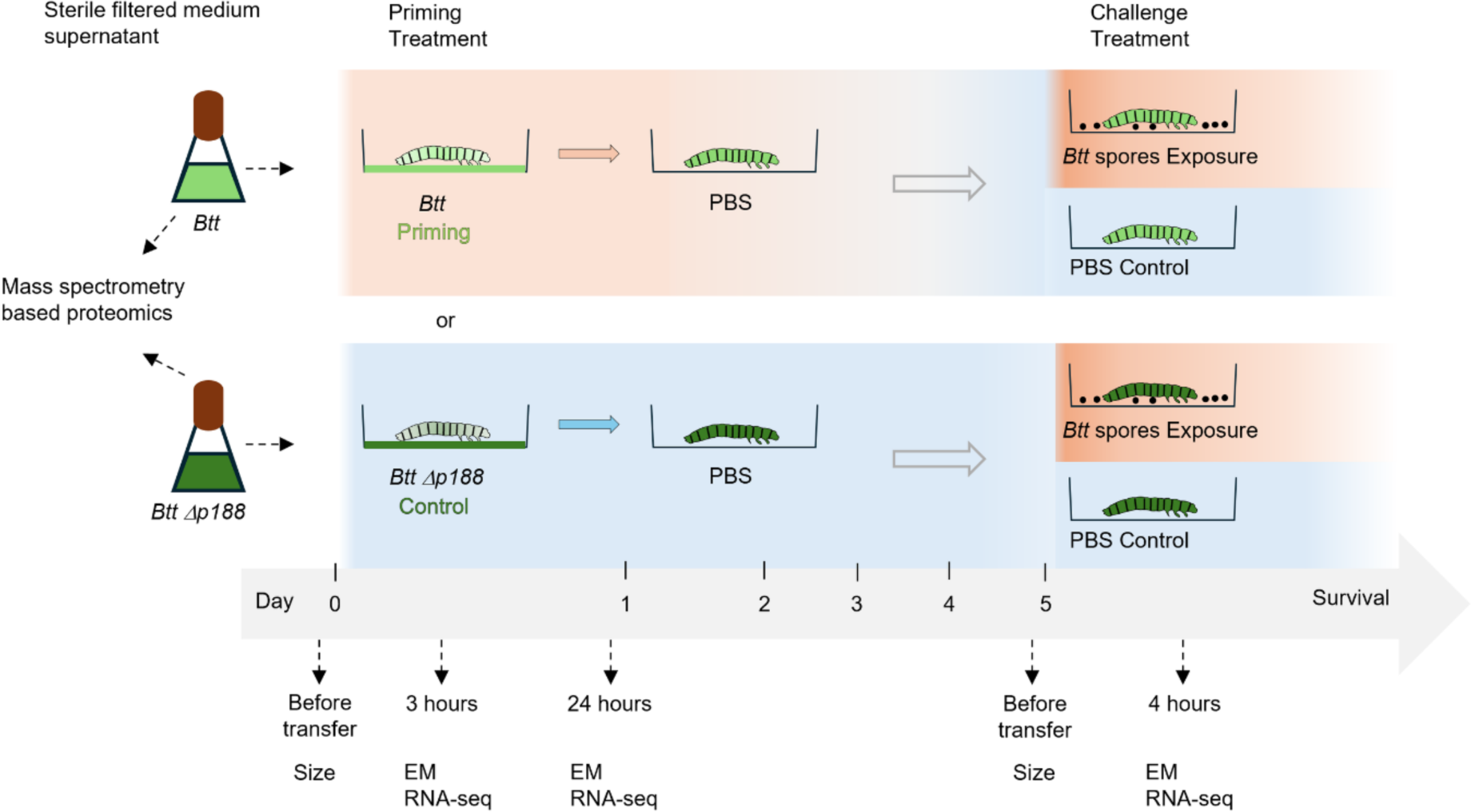
Study design. Before exposure to either priming- (*Btt*) or control (*Btt Δ p188*) supernatant flour mixes (Priming treatment) we measured the size of 14-day-old larvae. Early time-point gut samples were taken at 3 hours and 24 hours post-exposure for electron microscopy (EM) and RNA-sequencing (RNA-seq). The remaining larvae were transferred onto PBS flour mixes after 24 hours and left for four more days. Before transferring onto control (PBS)- or exposure (*Btt* spores) flour mixes (Challenge treatment), we measured the size of the same larvae as in the beginning. Four hours after the transfer onto the challenge diets, gut samples were taken for both, EM and RNA-seq again. The sterile-filtered spent culture media supernatants of *Btt* and *Btt Δ p188* that were used for the Priming treatment were also compared regarding their proteome composition with LC MS/MS.

### Preparation of diets

The spore cultures were prepared as described in Länger and Baur *et al.* (2023) [32] with some modifications. In short, either 50 mL (Priming treatment) or 400 mL (Exposure treatment) *Bt* growth medium was supplemented with 250 µL or 2 mL of a salt solution and 62.5 µL or 250 µL of 1 M CaCl_2_. Lastly, 1 mL or 5 mL of an overnight *B. thuringiensis* culture (*Btt* or *Btt Δ p188*) was added.

For producing the priming treatments, the sporulated cultures were centrifuged after seven days twice at 2604 rcf for 15 min. The resulting supernatants were filtered through a 0.45 µm filter followed by a 0.22 µm filter to remove bacterial spores and vegetative cells. Next, the filtered supernatants were mixed with 0.15 g flour/mL of supernatant, and 10 µL of each mix was pipetted into the wells of a 96-well flat bottom plate (Greiner). Finally, the plates were put to 30°C overnight for drying.

To produce the exposure treatment, the sporulated *Btt* cultures were centrifuged as described above. Between the first and second centrifugation the supernatants were discarded, and the spore pellets were washed in 15 mL PBS. The final spore pellets were resuspended in 5 mL PBS, the spores were counted in a Thoma counting chamber (depth 0.02 mm, square area 1/400 mm^2^) and the concentrations were adjusted to 1×10^10 spores/mL of PBS. Finally, 0.15 g flour was added to 1 mL of the spore/PBS mix and 10 µL of this final mix was pipetted into the wells of a 96-well flat bottom plate. The plates were put to 30°C overnight for drying.

#### Priming and challenge experiments

Priming and challenge of *T. castaneum* larvae was performed as described in [28]. In short, on day 15 after oviposition larvae were individualized into the wells of the control or priming diets (*Btt Δ p188* or *Btt*) or medium control flour diets (0.15 g flour/mL *B. thuringiensis* medium). After 24 hours the larvae were transferred onto PBS diets (0.15 g flour/mL PBS) where they were left for four days at standard conditions (for an overview see Fig 1). Finally, the larvae were transferred onto the exposure (*Btt* spores) or control (PBS) diets, and their survival was recorded daily for five days.

### Proteomics

To correlate differences in the ability to induce immune priming to differences in the proteomes of the spent growth media supernatants of *Btt* and *Btt Δ p188* cultures, we used four replicate cultures of both bacterial strains (supernatants prepared as in “Preparation of priming and infection diets“). Proteins from the supernatant were directly processed for MS analysis following the SP3 protocol, using TCEP (5 mM) and CAA (14 mM) for reduction and alkylation [33]. Proteins were bound to beads by the addition of 100 µL EtOH and washed twice with 80% EtOH. They were digested in 50 mM TEAB, pH 8.5 at 37 °C. After the overnight digestion, peptides were dried by vacuum centrifugation and reconstituted in 0.5% TFA and 2% acetonitrile for LC-MS/MS analysis. An EASY-nLC 1200 (Thermo Fisher) coupled to an Exploris 480 mass spectrometer (Thermo Fisher) was used for LC-MS/MS analyses. The peptides were separated on 20 cm frit-less silica emitters (CoAnn Technologies, 0.75 µm inner diameter), packed in-house with reversed-phase ReproSil-Pur C_18_ AQ 1.9 µm resin (Dr. Maisch) and the column was constantly kept at 50 C. Mass spectra were acquired in data-dependent acquisition mode as described in Sindlinger *et al*. [34]. The MaxQuant software version 2.0.3.0 [35] was used to process the raw data and standard settings were used. The LFQ count was 1. The resulting MS/MS spectra were assigned to a custom *Btt* proteome assembly (*Btt* genome kindly provided by Dr. Heiko Liesegang, Institute of Microbiology and Genetics, Georg-August University of Göttingen, unpublished) with default settings and match between runs. Label-free quantification (LFQ) and intensity based absolute quantification (iBAQ) options were enabled.

### Larval growth

To measure larval growth between priming and challenge time points, larvae were randomly selected before the transfer onto priming or control diets 14 days after oviposition and put onto petri dishes for area measurement. We measured the larval area again in the same way directly before the challenge treatment. The experiment was repeated in 3 blocks, resulting in 132 primed larvae and 135 control larvae being measured. To measure the area of the larvae, pictures were taken with an Olympus SZX12 microscope and saved as BigTIFF files. The CellSens Standard software was used to individually measure area in mm^2^. A small number of larvae, that had been exposed to priming or control diets, decreased in size, likely due to recent molting.

### Gut dissection

For both, electron microscopy (EM) and mRNA sequencing (RNA-seq), guts of larvae were dissected under sterile conditions after the larvae were immobilized on ice. To obtain guts for EM, larvae were placed into droplets of 15 µL cold PBS and the first and last segments were removed with a scalpel. The guts were then pulled out with forceps and put into 5 mL of fixing solution (see Method section Electron microscopy). Per timepoint and treatment, five guts were dissected for EM. For the guts used for RNA-seq 10 µL RNA-later (Thermo Fisher) was used instead of PBS. For each sample 7 guts were pooled in an 1.5 mL reaction tube containing 100 µL RNA-later on ice. Finally, the access RNA-later was removed, the samples were shock frozen in liquid nitrogen and stored at −80°. Per timepoint and treatment three replicates were used.

### Transmission electron microscopy

Dissected guts were fixed with 4% formaldehyde (Polysciences), 1% glutaraldehyde (Polysciences) in 0.1 M cacodylate buffer (Polysciences) pH 7.4 for 30 min and after a change for 3 hours at room temperature. The samples were stored at 4 °C until further processing. After fixation the samples were washed three times with 0.1 M cacodylate buffer (pH 7.4), post-fixed in 1% osmiumtetroxide (Polysciences) in 0.1 M cacodylate buffer at room temperature for one hour and washed with 0.1 M cacodylate buffer. Samples were gradually dehydrated in increasing concentrations (50 %, 70 %, 90 %, 96 %, 99.8 %) of ethanol (Roth) for 30 min at each step at 4 °C. After incubation with 100 % propylene oxide (Serva) twice for 10 min, the samples were embedded under vacuum by subsequent incubation. Epoxy resin (Serva) mixed with propylene oxide (1:1, 2:1 each step 4°C overnight), epoxy resin 100 % (2x 2 hours RT) and polymerized in fresh epoxy resin at 60 °C for 72 hours. Ultrathin sections (60-nm) were generated using a Leica EM UC7 ultramicrotome. Sections were poststained with 4 % uranyl acetate (Polysciences) in 25% ethanol and ‘Reynold’s lead citrate’. Samples were analyzed at 80 kV on a FEI-Tecnai 12 electron microscope (FEI). Images of selected areas were documented with Veleta 4k CCD camera (emsis).

### RNA-seq

To isolate total RNA, dissected gut samples were crushed in 1 mL of TRIzol (Thermo Fisher), sonicated, and centrifuged at 13000 RCF for 5 min. at 4 °C. After a chloroform wash and another round of centrifugation at 10500 RCF for 15 min. at 4 °C, the aqueous phase was collected in EtOH and RNA lysis buffer from the SV Total RNA Isolation kit (Promega), and the manual instruction was followed. The RNA was eluted in 70 µL of RNAse-free water. 15 µL of eluted RNA (∼ 80 ng/µL) was sequenced by Novogene (Cambridge, UK) using the Illumina NovaSeq PE150 platform (Paired-end, 150 bp read length). Raw reads were assessed using FastQC software (version 0.11.2) [36]. Adapters were removed and reads were filtered based on criteria such as adapter contamination, nucleotide uncertainty (if > 10% of a read), or low-quality nucleotides (if Base Quality < 5 in > 50% of a read). The resulting clean reads were mapped against the *T. castaneum* genome (Tcas5.2.54, http://ftp.ensemblgenomes.org/pub/metazoa/release-54/gtf/tribolium_castaneum/) using hisat2 (version 2.0.5) [37], assembled into transcripts or genes with Stringtie (version 1.3.3b) [38] and counted with featureCounts (version 1.5.0-p3) [39].

### Statistical Analysis

All statistical analyses were performed with R (version 4.3.1) [40] implemented in Rstudio (version 2023.06.1+524). Visualization of data was carried out using the R package “ggplot2” (version 3.4.4) [41].

For the proteome data, log_2_ transformation of LFQ intensities was performed. Only protein groups quantified in at least 3 of 4 replicates in at least one treatment supernatant (*Btt* or *Btt Δ p188*) were considered for the downstream analysis and missing LFQ values were imputed based on quantile regression using the package “imputeLCMD” (version 2.1). The R package “limma” (version 3.58.1) was used to statistically test for differential expression [42].

To statistically assess the influence of priming treatment on the growth of the larvae, measured area was modeled as the response variable explained by time point and treatment (priming or control) as explanatory variables and both, experimental blocks and individuals as random factors using the “lmer” function from the R package “lme4” (version 1.1-35.1) [43]. The model was as follows: Size ∼ Treatment*Time + (1|Block/Individual). To assess the treatment effect the full model was compared with a reduced model (Size ∼ 1 + (1|Block/Individual)) using the “anova” function.

For the survival analysis of the larval growth experiment, we applied a Cox proportional hazards model with one random factor using the “coxme” function from the “coxme” package (version 2.2-18.1) [44]. The model was as follows: (Surv(Day, Event) ∼ Treatment +(1|Replicate)). To assess the treatment effect, the full model was compared to a reduced model: “coxme” (Surv(Day, Event) ∼ 1 +(1|Experiment), data), using the “ANOVA” function. The medium control was set as the reference. For the survival analysis of the RNA-seq and EM experiments, we applied the “coxph” function from the “survival” package (version 3.5-7) without adding a random factor: (Surv(Day, Event) ∼ Treatment). In all cases, treatment (medium, *Btt* or *Btt Δ p188*) was set as the fixed factor, and proportionality of the fixed effect was assessed with the cox.zph() function from the “survival” package. Additionally, the survival curves were plotted using the “survfit” function from the “survival” package [45] and “ggsurvplot” from “ggplot2”.

For the RNA-sequencing data, differential gene expression analysis was performed with DESeq2 (version 1.20.0) [46], on genes with raw counts greater than 2 in the comparison groups (priming vs. control after 3 hours, 24 hours, 5 days, 5 days + *Btt*). Genes with a log2 fold change > 0.5 or < −0.5 and a False Discovery Rate (FDR, Benjamini-Hochberg) adjusted p-value < 0.05 were considered significantly up-regulated or down-regulated, respectively. GO enrichment analysis was performed using the R package clusterProfiler (version 4.10.0) [47] with a threshold of FDR adjusted p < 0.05. Weighted Gene Co-expression Network Analysis was carried out using the R package WGCNA (version 1.72-5) [48]. Initially, samples were checked for outliers (hierarchical clustering upon distance matrix computation and PCA), genes with a raw count lower than 10 in 50% of the samples were filtered out, and the remaining reads were variance stabilized using the “vst()” function (DESeq2). A soft thresholding power for gene correlations of 14 was chosen based on high-scale freeness and low mean connectivity. Both network type and TOMtype were set to “signed” to cluster genes that are either co-up- or co-down expressed. To statistically compare the module eigengene values of the different modules between groups, the R package limma (version 3.58.1) was used. Modules with significantly different module eigengene values (FDR adjusted p < 0.05) were exported to Cytoscape (version 3.10.1) [49] and visualized with the plug-in ClueGO (version 2.5.10) [50]. Only significantly enriched terms (FDR adjusted p < 0.05) were visualized and terms were grouped based on the number of shared genes (Kappa score = 0.4).

## Results

### *T. castaneum* oral immune priming response in the gut peaks after 24 hours and resembles an infected state

As little is known about the mechanisms of oral immune priming in the insect gut, we here used the *T. castaneum – B. thuringiensis tenebrionis* host-pathogen model to provide insights into this phenomenon. The survival data derived from the present experiments confirm oral immune priming in *T. castaneum:* larvae showed improved survival of *Btt* infection when they were previously exposed to priming diets derived from *Btt* culture supernatants (Estimate = 0.57, p < 0.001), but not when exposed to *Btt Δ p188* control diets (Estimate = 1.13, p = 0.34) (each compared to *Bt* medium diets, Fig 2A,B, S1 Fig).

**Fig 2:**
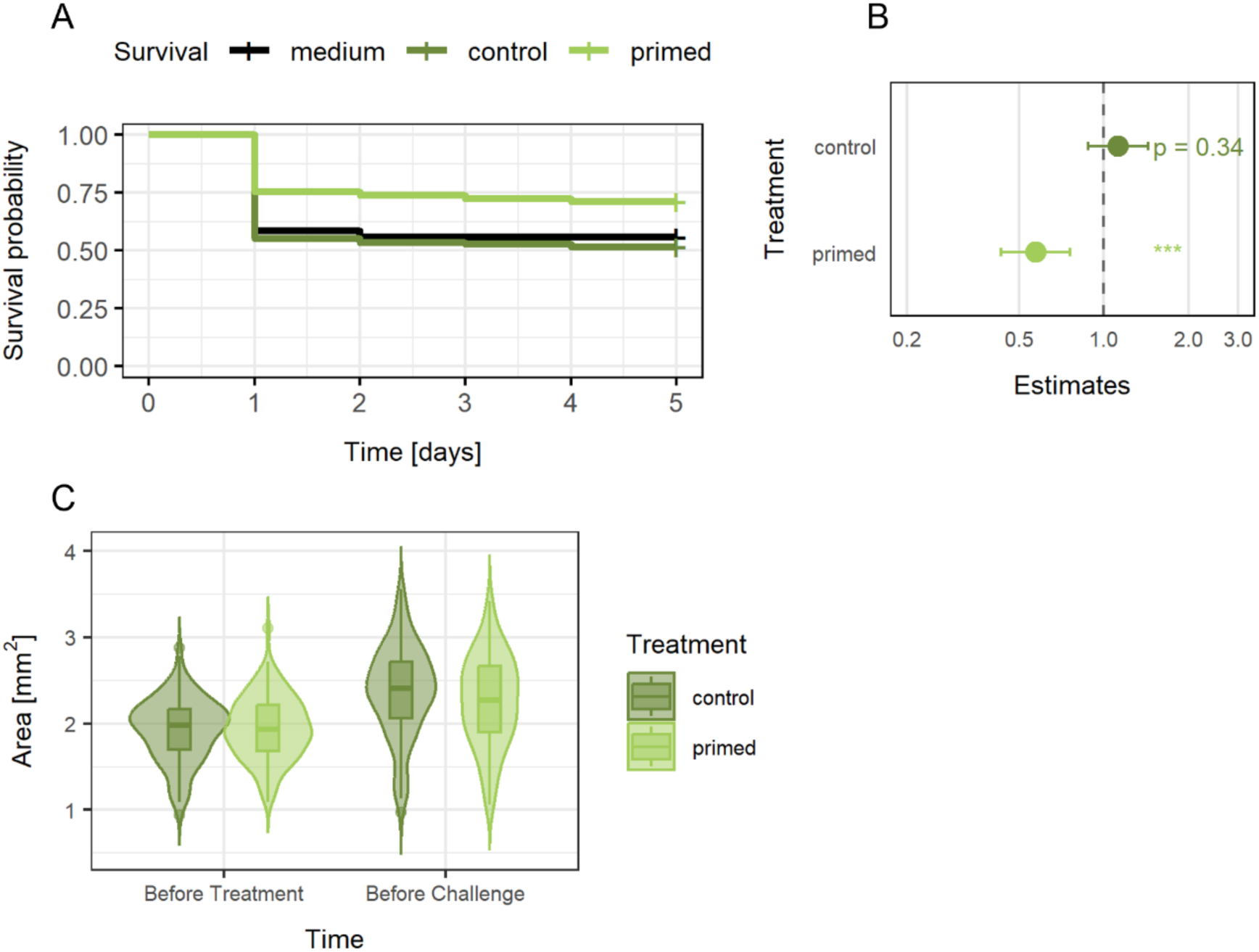
Infection survival and growth of *T. castaneum* larvae after priming. **A** Kaplan Meier curves displaying the survival of *T. castaneum* larvae exposed to either *Bt* medium diets (n =495), control diets (n =500), or priming diets (n =501) upon exposure to infectious *Btt* spores. **B** Forest plot representing the estimated fixed-effects coefficients of primed and control compared to medium (black, dashed line a 1.0). **C** Growth of *T. castaneum* larvae exposed to either priming (n = 132 larvae) or control (n = 135 larvae) diets. Larval size is given as their body area. The raw data for survival and growth can be found in S1 Table.

The increased survival probability after priming came at the cost of reduced growth in the treated individuals, which can be an indicator of slower development to pupal and adult stages [9]. During the time between priming and challenge exposure, larvae, which were initially of the same size (Fig. 2C, Estimate = 0.02038, p = 0.6586) grew significantly less if they received the priming treatment compared to the control (Estimate = −0.13439, p < 0.001).

To analyze gene expression patterns in gut tissues from larvae that were either exposed to priming or control diets we combined two approaches: DESeq2 was used to compare each gene’s expression between priming and control at each time point [45], and WGCNA was used to construct gene co-expression networks [47]. The number of differentially expressed genes (DEGs) between primed and control larval gut tissues varied over time (Fig 3A), peaking at 24 h post-exposure with approximately 5 – 10 times as many DEGs compared to the other time points. In unchallenged larvae, more genes were down-regulated in primed larval gut tissues after five days, whereas upon challenge with life *Btt* most DEGs were up-regulated in primed larval gut tissues (Fig 3A). We also used the normalized gene expression of all DEGs identified between treatments across time points and performed hierarchical clustering, as well as principal component analysis (Fig 3B, S2 Fig). The samples clustered well by their group (time and treatment, Fig 3B). Moreover, samples that were exposed to *Btt* spores (both primed and control) clustered together. However, the primed samples after three and 24 hours clustered closest with one another and the *Btt* exposed samples, whereas the primed samples after 5 days clustered closest with the control samples. This shows, that early after priming and exposure to *Btt* spores many genes are similarly strongly expressed. When comparing the identity of DEGs between priming and control across the different time points, there was little overlap indicating that the priming response generally is dynamic (Fig 3C, D).

**Fig 3:**
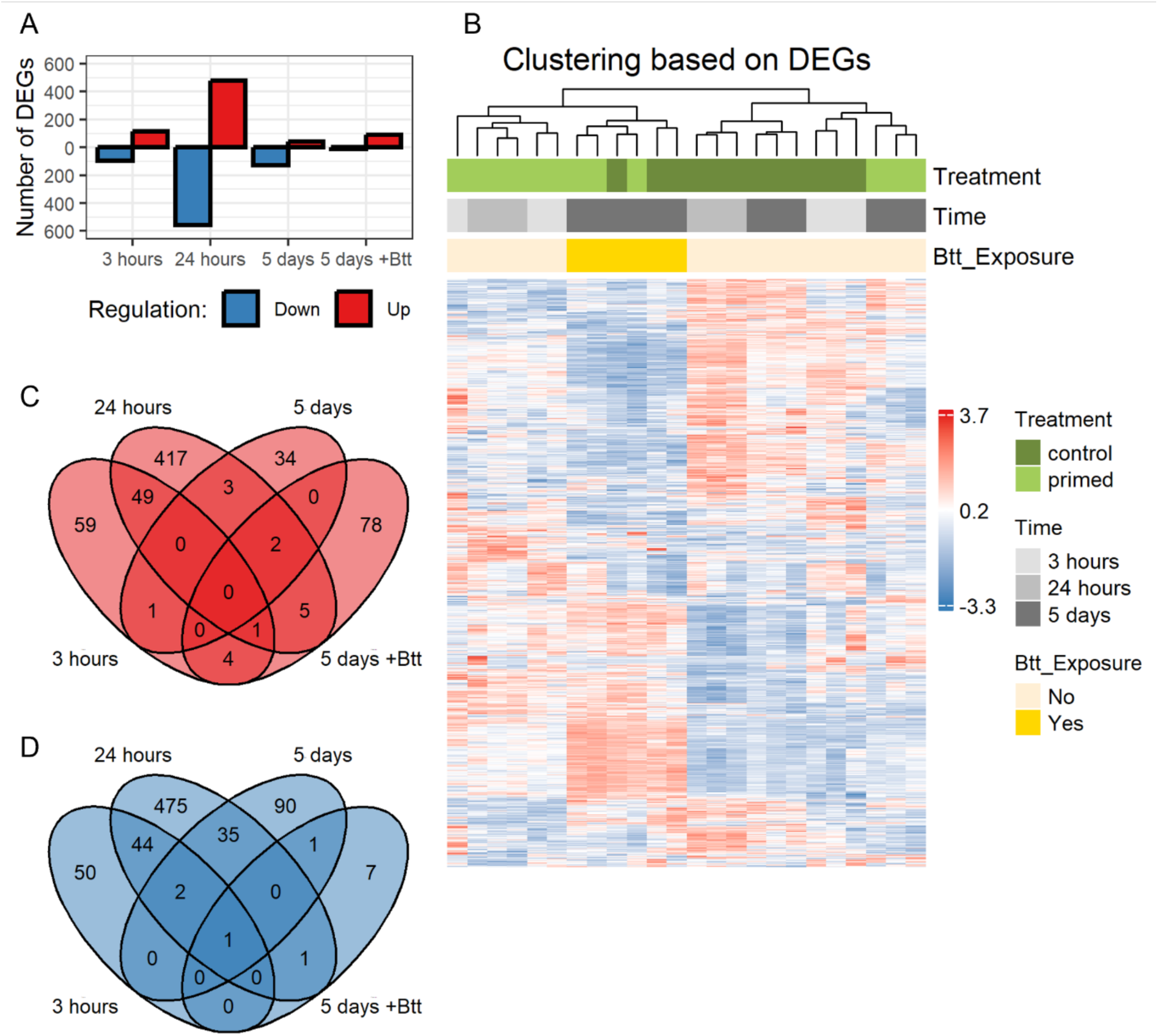
Overview of the results from the DESeq2 analysis. **A** Number of differentially expressed genes in guts from *T. castaneum* larvae exposed to priming *vs.* control diets (Down = lower expressed in primed guts, Up = higher expressed in primed guts), three hours, 24 hours and 5 days after priming without and with challenge (+Btt), analyzed with DESeq2 (Differentially expressed: Down in primed guts = Log2FC < −0.5, FDR adjusted p < 0.05, Up in primed guts = Log2FC > 0.5, FDR adjusted p < 0.05). **B** Heatmap representing the hierarchical clustering of the normalized gene expression for DEGs across all samples. Rows represent all DEGs identified in this study at all timepoints between primed and control larvae. Columns represent individual triplicate samples grouped by treatment, Time and exposure to infectious *Btt* spores (No = not exposed, Yes = exposed). Colors of the cells indicate the correlation of gene expression for a gene (row), in a sample (column). **C and D** Venn diagram for significantly up- (**C**) and down-regulated (**D**) genes between primed and control larval gut tissues at the different timepoints (three hours, 24 hours, 5 days, 5 days exposed to *Btt* spores). Overlapping areas represent shared DEGs between timepoints. The raw count data from the RNA-sequencing run can be found in S2 table.

After filtering out genes with low expression levels across samples, 10,814 genes of the total 17,052 genes remained for WGCNA. The WGCNA analysis identified 25 modules, with module 1 containing 2,262 genes and module 25 containing 36 genes (S3 Fig). Similarly, to the differential gene expression analysis, *Btt* exposed samples from control and priming treatment clustered well together upon Pearson correlation of the module eigengene values with the samples grouped by treatment and time (S4 Fig). We then went on and statistically compared the module eigengene (ME) values at each timepoint between primed and control samples. Only the 24-hour and 5-day time points displayed statistically different ME values. After 24 hours, 7 modules displayed statistically different ME values (FDR adjusted p < 0.05) between primed and control samples (Modules 1, 2, 8, 11, 13, 14, 17, Fig 4). Module 8 was excluded as no GO terms could be identified in this module. The gene ontology (GO) analysis of key modules revealed that biological processes associated with protein synthesis and folding are upregulated in primed larval guts after 24 hours compared to controls, suggesting an increased cellular stress response (Fig 4A, Module 1). Additionally, priming was associated with elevated expression of genes involved in cell cycle regulation and nuclear DNA-related processes, further indicating an increased stress state (Fig 4A, Module 13). Conversely, Modules 2, 11, 14, and 17 comprised genes downregulated in primed larvae relative to controls (Fig 4B). Processes such as lipid catabolism and gene expression were reduced, which, in conjunction with the upregulation of protein folding genes, may indicate endoplasmic reticulum (ER) stress. Downregulation of genes involved in cellular signaling, including the BMP signaling pathway, paired with upregulation of cell cycle genes, which might suggest shifts in cell differentiation. Notably, at the 5-day mark, Modules 14 and 17 also exhibited significant differences in module eigengene (ME) values between primed and control guts (Fig 4B). These WGCNA findings underscore the potential for priming-induced stress, prompting us to combine transcriptomic insights with transmission electron microscopy (TEM) to investigate ultrastructural changes associated with gene expression dynamics in response to priming.

**Fig 4.**
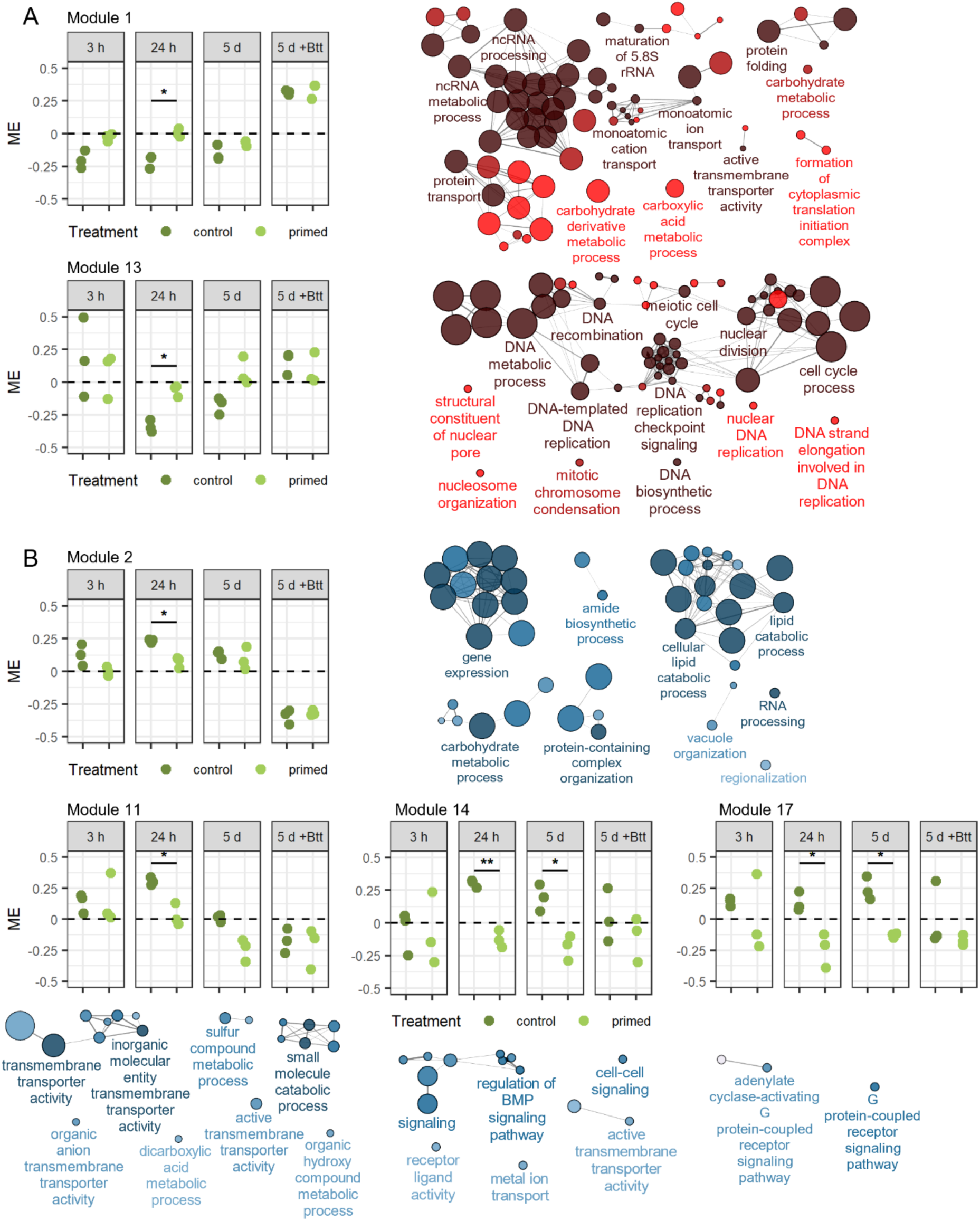
WGCNA modules with a significantly different module eigengene value (FDR adjusted p < 0.05) between larvae exposed to priming or control diets. The dot plots display module eigengene values (y-axis) vs samples grouped by timepoint (h = hours, d = days) and priming treatment (x-axis). Asterisks display significance levels (* = p <=0.05, ** = p <= 0.01). The networks represent Biological Processes contained in the modules identified with ClueGO. Only significantly enriched terms are included (Holm-Bonferroni adjusted p < 0.05). Each node represents a GO term. Node labels are based on the most significant GO term of a group of interconnected GO terms. Node size represents the number of genes mapped to the node. Node color reflects significance level (Dark = highly significant, bright = less significant). Edges connect nodes and define groups based on shared genes (Kappa >= 0.4). **A** Modules 1 and 13 show higher module eigengene values for primed compared to control. **B** Modules 2, 11, 14, and 17, show lower module eigengene values for primed compared to control. WGCNA analysis is based on the data found in S2 and S3 Table.

### The early priming response is characterized by damage-associated stress responses and an up-regulation of immune genes

As early as three hours after the exposure to the priming diets, the electron micrographs revealed a tattered peritrophic matrix (compare Fig 5A and 5D, arrowheads) and a higher abundance of apoptotic cells in the gut of primed larvae (compare Fig 5C and Fig 5F). We also observed an increase of membrane vesicles in the gut lumen after priming (compare Fig 5B and 5E). Fitting these pathology observations in the gut of primed larvae, many DEGs at this time point are related to metabolic processes (Fig 5G). Up-regulated genes in primed larvae fell into the category of serine-type endopeptidases (S4 Table: 3 h: Category = “Proteases”), but also a ceramide synthase and genes encoding for proteins involved in intracellular protein degradation were significantly higher expressed (S4 Table: 3 h: Category = “Lipid metabolism” and “Proteases”). Most down-regulated genes fell into categories of lipid and carbohydrate catabolic processes (Fig 5G, S4 Table: 3 h: Category = “Lipid metabolism” and “Glycosidases”). None of the modules derived from the WGCNA analysis displayed significantly different ME values three hours after priming, but most up-regulated genes identified by DESeq2 clustered into module 1, and most down-regulated genes clustered into module 2.

**Fig 5:**
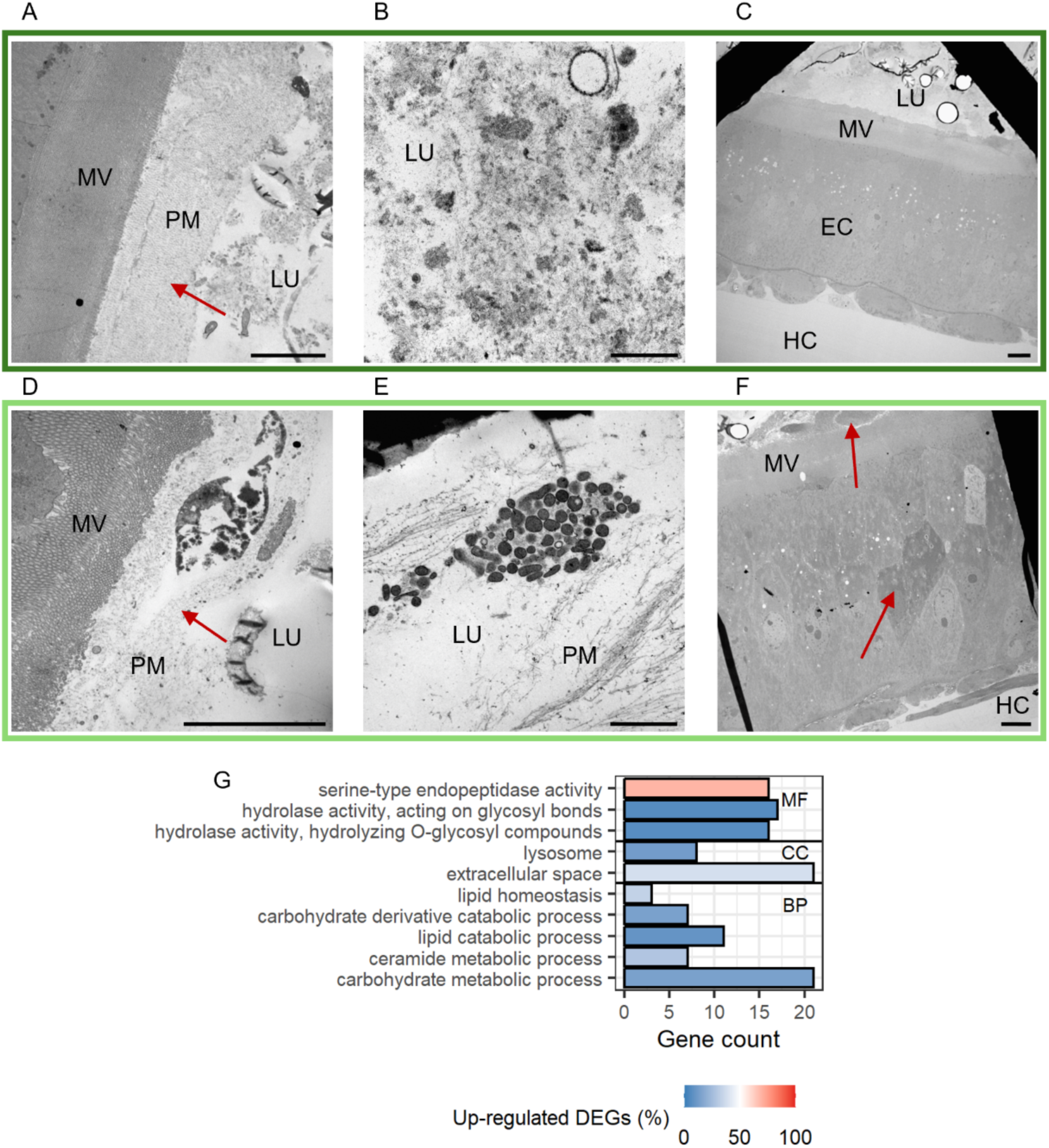
Differences between larvae exposed to control (*Btt Δ p188*) and priming (*Btt*) diets 3 hours after exposure. **A – F** Electron micrographs of control larvae (A-C) and primed larvae guts (D-F). Micrographs were chosen based on similar position and depth of sections between treatments. LU = gut lumen, MV = microvilli, PM = peritrophic matrix, EC = epithelial cell, HC = hemocoel. Red arrows: A, D = highlighting differences in peritrophic matrix, F = apoptotic cells. Scale bars: A = 5 µm, B = 1 µm, C = 5 µm, D = 5 µm, E = 1 µm, F = 5 µm. **G** GO term enrichment analysis for differentially expressed genes (DESeq2) between primed and control larval guts. Only GO terms with a significant enrichment are displayed (FDR adjusted p < 0.05). MF = Molecular Function, CC = Cellular Compartment, BP = Biological Process. The x-axis shows the number of genes that belong to a GO term, the color gradient indicates the ratio of down- (blue) and up-regulated (red) genes identified with DESeq2.

In line with the peak in the number of DEGs after 24 hours, we detected stronger differences in the gut ultrastructure between primed and control larvae at this time point. In primed larvae, the tattered peritrophic matrix and increased apoptosis detected after three hours persisted. Still, additional structures in the gut lumen and in between microvilli appeared, which seem to be membrane residues originating from gut epithelial cells (compare Fig 6A and 6D, arrowheads). Both WGCNA and DESeq2 revealed down-regulated genes in primed larval gut tissue that clustered into GO categories of lipid and carbohydrate catabolic processes (Fig 4B: Module 2, Fig 6G, S4 Table: 24 h: Category = “Lipid metabolism” and “Glycosidases”), which coincides with the disturbance of cellular membranes and the PM on the ultrastructure level. At the same time, genes encoding for chitin synthase and ceramide synthase were up-regulated, possibly counteracting the degradation of cellular membranes and the PM. Also, mitochondria in the epithelial cells of primed larval guts showed signs of stress, such as elongated shapes and shrinkage (compare Fig 6B and E, arrowheads). Additionally, we observed an increased number of autolysosomes, presumably due to the increased degeneration of mitochondria (compare Fig 6C and 6F). This stress on the ultrastructure level within the epithelial cells was also reflected by a high number of genes that clustered into GO terms that involve the synthesis, modification, transport, and degradation of proteins, especially in the endoplasmic reticulum, which was again indicated by both DESeq2 and WGCNA (Fig 4A: Module 1, Fig 5G, S4 Table: 24 h: Category = “ER stress”). We also found the GO term “defense response to bacterium” to be significantly enriched in primed larval guts at this time point (Fig 6G). Within this category are two gene copies of *Attacin*, *Coleoptericin,* and *Defensin*, respectively, as well as a gene encoding for an inducible metalloprotease inhibitor (S4 Table: 24 h: Category = “Immune”). Also, two genes encoding for peptidoglycan recognition receptors were differentially regulated. Further significant modules identified by WGCNA with higher gene expression profiles in primed larval gut tissues contained genes that participate in the recombination, replication, and repair of DNA and genes associated with cell cycle processes in the nucleus (Fig 4A: Module 13). Modules that were associated with a lower gene expression profile contained genes involved in transmembrane transporter activities and cell-to-cell signaling (Fig 4B: Modules 11, 14, and 17).

**Fig 6:**
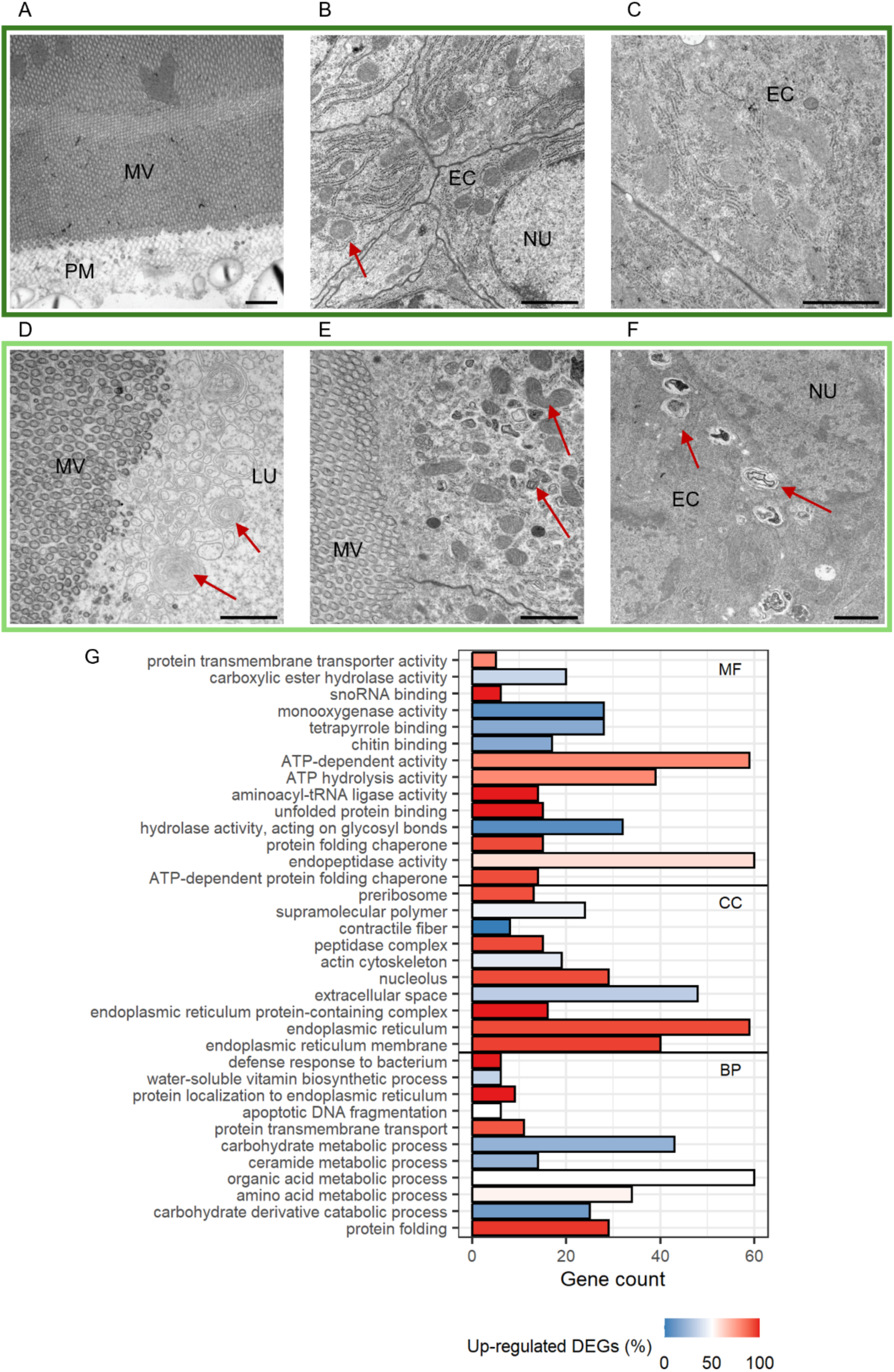
Differences between larvae exposed to control (*Btt Δp188*) and priming (*Btt*) diets 24 hours after exposure. **A – F** Electron micrographs of control larvae (A-C) and primed larvae guts (D-F). Micrographs were chosen based on similar position and depth of sections between treatments. LU = gut lumen, MV = microvilli, PM = peritrophic matrix, EC = epithelial cell, NU = nucleus. Red arrows: A, D = “normal” (A) and “stressed” (D) mitochondria, E = autolysosomes, F = putative membrane residues originating from epithelial cells. Scale bars: A - F = 5 µm. **G** GO term enrichment analysis for differentially expressed genes (DESeq2) between primed and control larval gut tissues. For details, see Fig 5.

### Reduced gene expression differences after 5 days, while stress signs at the ultrastructural level persist

After 5 days, we still observed structural differences in the gut of primed larvae, such as the above-mentioned unusual shapes of mitochondria or the increased abundance of autolysosomes (compare Fig 7A and 7D). Additionally, we detected the occurrence of signs of stress in the ER of epithelial cells, namely the degeneration of ribosomes and even the degradation of the ER in primed larvae (Fig 7D). The most prominent change, however, was the microvilli (MV) swelling in primed larval guts (compare Fig 7B, E, C, F). These swellings occurred towards the tip of the MV and finally led to the extrusion of MV debris. Genes that were down-regulated in primed larvae at this time point encoded, among others, for proteins that are part of the cytoskeletal architecture or are involved in detoxification processes (Fig 7G, S4 Table: 5d: Category = “Cytoskeletal”, “Monooxygenases”). After 24 hours, genes encoding for transmembrane transporter activity were down-regulated as well as revealed by WGCNA and DESeq2 (Fig 4B: Modules 14 and 17, Fig 7G, S4 Table: 5d: Category = “Transporters”).

**Fig 7:**
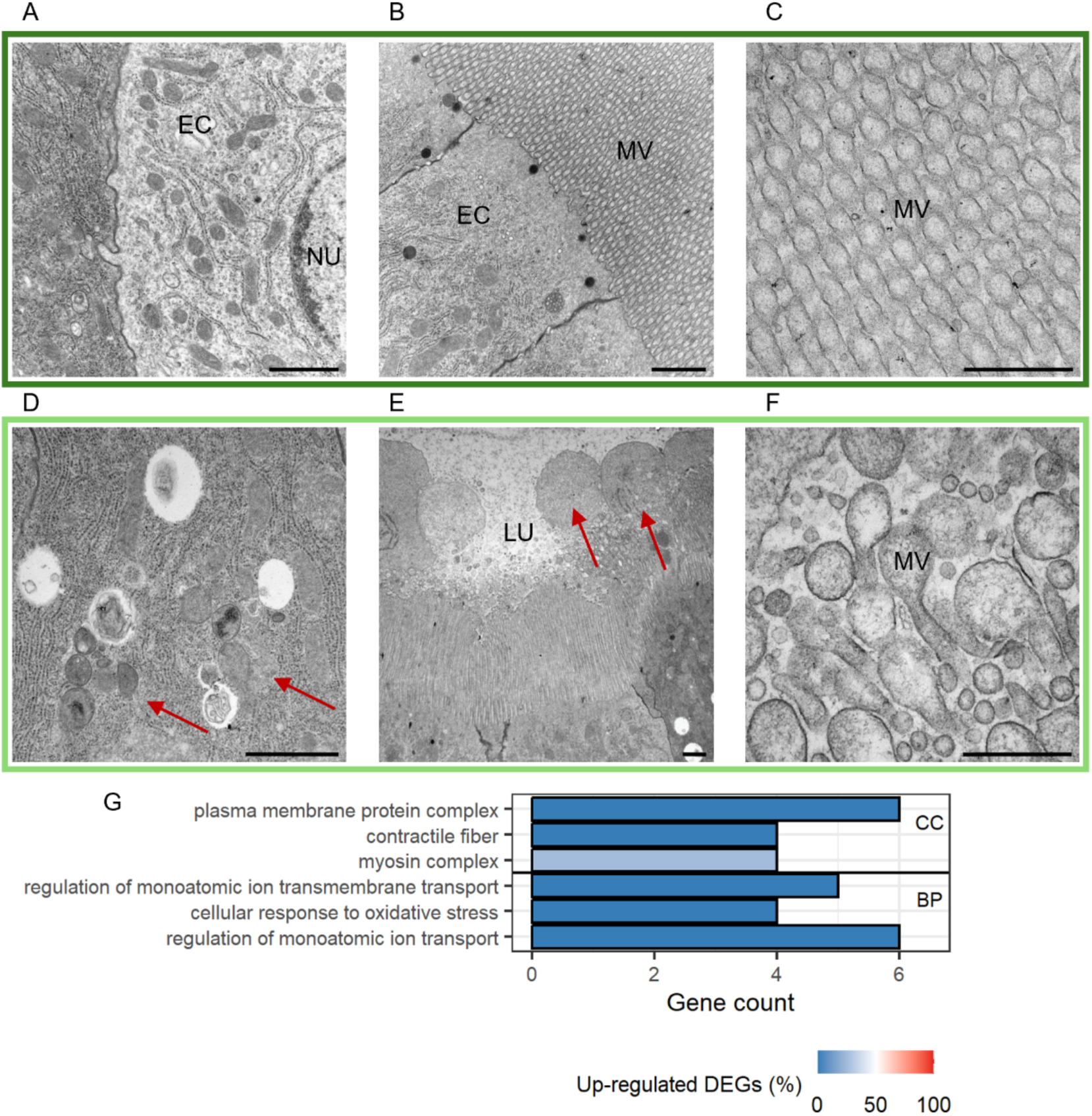
Differences between larvae that had been exposed to control (*Btt Δ p188*) and priming (*Btt*) diets 5 days after exposure. **A – F** Electron micrographs of control larvae (A-C) and primed larvae guts (D-F). Micrographs were chosen based on similar position and depth of sections between treatments. LU = gut lumen, MV = microvilli, EC = epithelial cell, NU = nucleus. Red arrows: D = stressed endoplasmic reticulum, E = swollen microvilli. Scale bars: A = 1 µm, B = 1 µm, C = 500 nm, D = 1 µm, E = 1 µm, F = 500 nm. **G** GO term enrichment analysis for differentially expressed genes (DESeq2) between primed and control larval gut tissues. For details, see Fig 5.

### Upon exposure to life *Btt*, primed guts show higher expression of genes involved in detoxification processes

Five days after exposure to the priming or control diets, the guts of larvae that were exposed to *Btt* spores showed signs of infection in all samples. Vegetative *Btt* cells (Fig 8A and D), lysosomal structures (Fig 8B), and intracellular vesicles (Fig 8E) were present in both control and primed larvae. In some gut sections, we detected an increased number of apoptotic cells (Fig 8C), whereas other sections displayed a stressed MV rim (Fig 8F). Primed and control larval guts hardly differed, as the biggest influence on gut integrity seemed to be the number of germinated *Btt* cells, which varied largely even within the treatment. In line with this, the number of DEGs between primed and control larvae was the lowest among the investigated time points (Fig 3A). The only significant GO term containing down-regulated genes was “beta-glucosidase activity”, including an acidic chitinase (Fig 8G). Genes that were up-regulated in primed larvae upon *Btt* exposure encoded for detoxification enzymes, which were down-regulated in non-exposed larvae at this timepoint, lipid binding proteins and proteins containing leucine-rich repeats (Fig 8G, S4 Table: 5d_E: Category = “Detoxification”, “Lipid binding”, “LRR”). Similar to three hours after priming, none of the modules derived from the WGCNA analysis displayed significantly different module eigengene values after the spore exposure treatment.

**Fig 8:**
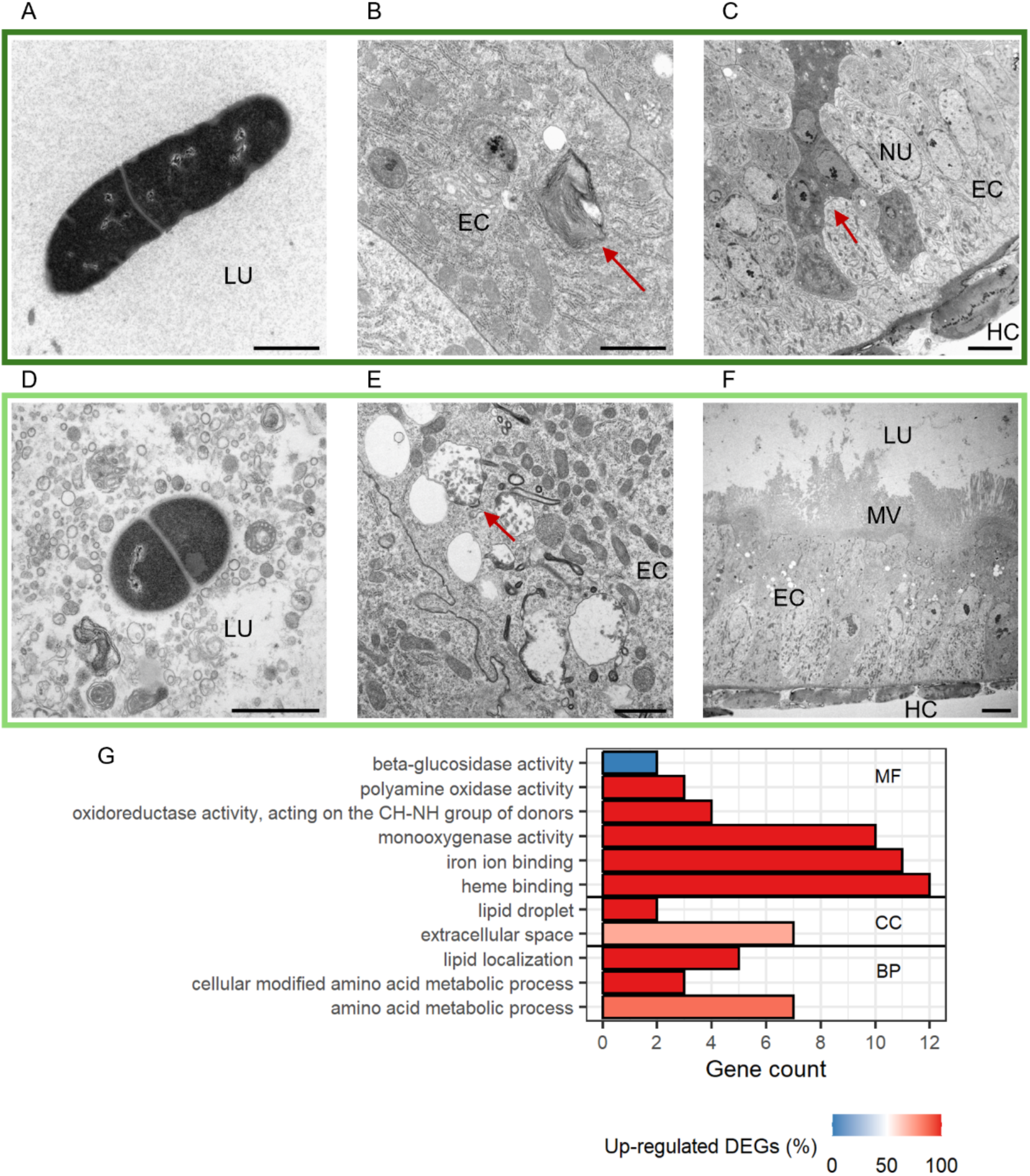
Differences between larvae that had been exposed to control (*Btt Δ p188*) and priming (*Btt*) diets 5 days after exposure and 4 hours after exposure to infectious *Btt* spores. **A – F** Electron micrographs of control larvae (A-C) and primed larvae guts (D-F). Micrographs were chosen based on similar position and depth of sections between treatments. LU = gut lumen, MV = microvilli, EC = epithelial cell, NU = nucleus, HC = hemocoel. Red arrows: B = lysosomal structure, C = apoptotic cells, E = intracellular vesicles. Scale bars: A = 1 µm, B = 1 µm, C = 5 µm, D = 1 µm, E = 1 µm, F = 5 µm. **G** GO term enrichment analysis for differentially expressed genes (DESeq2) between primed and control larval gut tissues. For details, see Fig 5.

### Priming-associated damage in the gut is likely caused by virulence factors present in the priming diet

When we compared the proteomes of priming (*Btt*) and control (*Btt Δ p188*) centrifuged, sterile filtered media supernatants, the only significantly differentially enriched protein group was a Sphingomyelinase C, which showed increased expression in the priming inducing *Btt* supernatant (S5A Fig). As expected, proteins that are encoded on the Cry-toxin carrying 188 kb plasmid were below detection limit in the *Btt Δ p188* control supernatants, where this plasmid was removed. Among those proteins were the crystal toxins Cry3Aa and Cry15Aa (S5A Table). Sphingomyelinase C is encoded on the same plasmid. The low expression level detected in 3 of 4 control media might be explained by another homologous Sphingomyelinase encoded on the chromosome and containing identical peptide fragments. Upon imputation of missing values, Cry15A, Cry3Aa and Sphingomyelinase C were among the few protein groups that were significantly higher expressed in priming spent culture media supernatants (Fig 9, S5B Fig, S5B and S5C Tables). Another virulence factor, Chitinase D, that was previously found to be differentially expressed in *Btt* compared to another *Bt* strain, was highly expressed in both, control and priming supernatants [32].

**Fig 9:**
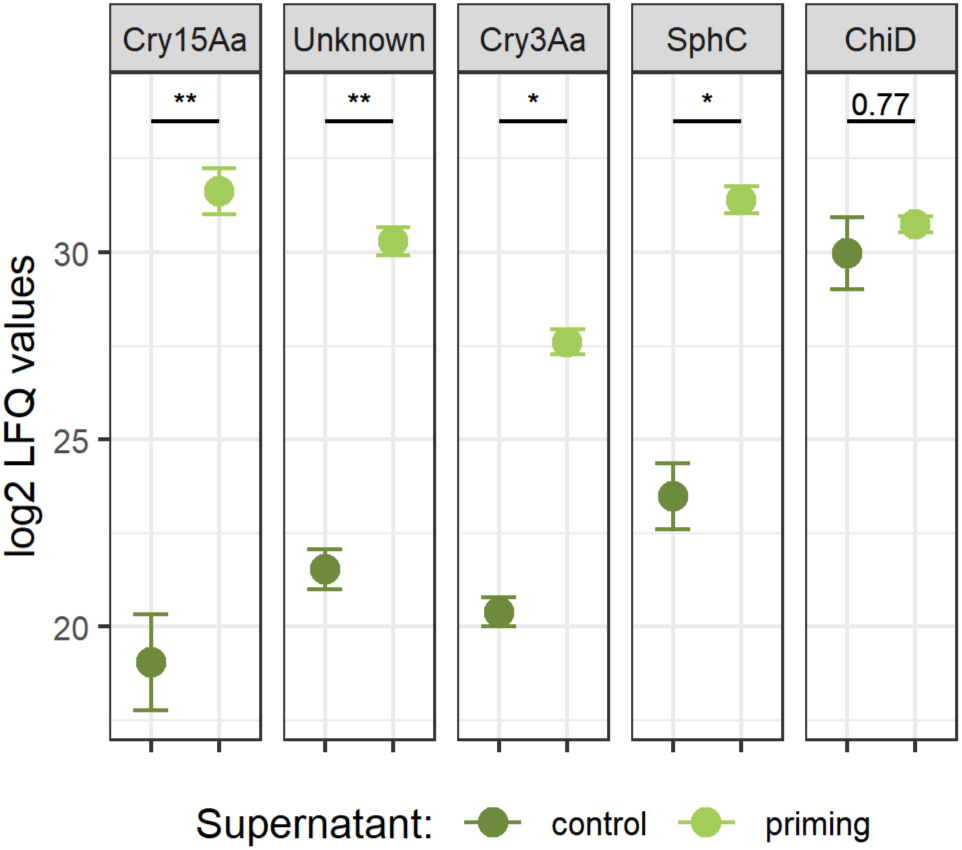
Exemplary proteome expression differences between priming (*Btt*) and control (*Btt Δ p188*) spent culture media supernatants. Depicted are the mean values of the normalized and log2-transformed label-free quantification values after imputation for missing values (n = 4). Error bars represent the standard error. ** = p value < 0.01, * = p value < 0.05. SphC = Sphingomyelinase C, ChiD = chitinase D.

## Discussion

To gain insights into the dynamic responses following oral immune priming in the insect gut, we combined electron microscopy and RNA-sequencing of guts of primed and control *T. castaneum* larvae at different timepoints and upon exposure to life *Btt* spores. Additionally, we used mass spectrometry to characterize the cues present in the priming inducing spent media supernatants of *Btt* cultures. Our results show a strong reaction in the primed gut associated with *Btt*-triggered tissue damage early after priming, including the up-regulation of immune genes, that diminishes over time but enables a targeted response upon re-encountering infective *Btt* spores.

The gut epithelium was strongly affected early after the exposure to the priming diet, as indicated by damage in the peritrophic matrix and an increase in apoptotic cells and autolysosomes after three hours (Fig 5D and 5F). After 24 hours, such damage was more pronounced; additionally, mitochondria showed signs of stress or degraded. Membrane residues were shed into the gut lumen (Fig 6A-F). Similar membrane and micro-vesicle shedding upon exposure to pore-forming toxins has been reported in HeLa cells and mice and seems to aid in the repair of damaged cellular membranes [51, 52]. Overall, the observed changes in gut ultrastructure in primed larvae at these early time points reflect the presence of virulence factors (e.g. Cry toxins, Chitinases, Sphingomyelinase C) in priming-inducing *Btt* culture supernatants and resemble natural *Bt* infections [29, 30]. Chitinases break down the chitinaceous peritrophic matrix, and Cry toxins, once dissolved and activated by midgut proteases, bind to receptors on midgut epithelial cells, ultimately leading to cell death [29, 53, 54]. Sphingomyelinases further damage cellular membranes by hydrolyzing sphingomyelin to ceramide and phosphorylcholine [55, 56]. In accordance with the increase of signs of damage after 24 hours, the number of differentially expressed genes (DEGs) between primed and control larval gut tissues also peaked at this time point. Among the DEGs were cysteine- and serine proteases, the main digestive enzymes in *T. castaneum* [57]. In a study in *Tenebrio molitor*, oral exposure to Cry3Aa led to a similar shift in the expression profiles of cysteine- and serine proteases [58]. Also, early work on the proteolytic processing of Cry3Aa by insect gut enzymes indicated an influence of the gut enzyme composition on Cry3Aa activity [59, 60]. After 24 hours, two serine protease inhibitors (serpins) were additionally upregulated in primed larval guts, suggesting active regulation, especially for these proteases. Serpins’ tight regulation of these proteases might also be associated with their roles in insect immunity [61]. Genes encoding for glycosyl hydrolases family proteins were downregulated at the early time points after priming. More than a third of the glycosyl hydrolases, which were downregulated after 24 hours, are involved in chitin degradation. This, together with the upregulation of chitin synthase, could indicate a strategy to suppress chitin catabolism in the presence of externally induced peritrophic matrix damage. Genes involved in lipid metabolic processes were differentially regulated after priming at the early timepoints as well. Ceramidases, sphingomyelinases, and lipases were mostly downregulated. Downregulation of acid- and neutral sphingomyelinases in mammalian cell lines aids regeneration upon exposure to pore-forming toxins [62]. The ceramide synthase *CerS6* gene and a gene encoding for a serine palmitoyltransferase, important in sphingomyelin synthesis, were upregulated in primed larvae at these early time points. Sphingomyelins are major constituents of cellular membranes but also play roles in endoplasmic reticulum (ER) homeostasis and immune signaling [63–66]. The presence of sphingomyelinase C in priming-inducing supernatant, together with the differential expression of sphingomyelin-associated genes and the occurrence of membrane residues in the gut lumen of primed larvae after 24 hours, suggests a disturbance of lipid homeostasis in these guts. In the intestines of cockroaches, it has been shown that ER stress and lipid homeostasis are connected, and that disturbance increases the expression of pro-inflammatory genes [67]. At 24 hours post priming, genes were up-regulated that are linked to such ER-associated stress. These included genes that encode for membrane-spanning proteins in the ER [68–71] and genes encoding for components of the cytosolic proteasome complex, necessary for the correct degradation of proteins [72]. Also, genes that are known to play important roles in the unfolded protein response (UPR), a cellular stress response related to ER stress, were up-regulated [73–77]. In *Aedes aegypti,* knockdown of UPR genes reduced the survival upon treatment with Cry11Aa toxin [78]. Most of these genes clustered into module 1 in our WGCNA analysis whose eigengene value was significantly higher in primed larvae than control larvae after 24 hours (Fig 4A).

Cell cycle and checkpoint signaling genes were also higher expressed in the primed gut after 24 hours compared to control guts (Fig 4A). Many genes involved in these processes have been reported to correlate with ER stress, and the influence of ER stress on DNA integrity, repair and epigenetic changes, is a topic of active research [79–81]. Further, differences in gene expression in the cell cycle could hint at divergent stem cell decisions upon priming, as observed upon pathogen exposure in Drosophila [82]. Unresolved ER stress leads to increased autophagy and cell death, signs we detected in the ultrastructure of primed larvae at these early time points. Generally, genes involved in gene expression were expressed lower in primed larvae (Fig 4B). In contrast, tRNA synthetase- and ribosome genes were up-regulated (Fig 6G), potentially reflecting an increased demand in protein synthesis while controlling for unbalanced gene expression [83]. Two genes encoding peptidoglycan recognition proteins (PGRP) were differentially regulated after 24 hours. PGRP-SA was up- and PGRP-LB down-regulated. The same direction of expression for these two PGRPs has previously been observed following septic infections in *T. castaneum* using the Gram-positive bacterium *Micrococcus luteus* [84]. Additionally, Gram-positive bacteria targeting antimicrobial peptides (AMPs) were up-regulated after 24 hours [85–87], indicating a strong immune reaction 24 hours after exposure to the priming diet. Interestingly, genes encoding for monooxygenases, a group of enzymes that are known to be involved in detoxification processes and are overexpressed in some *Bt*-resistant insects [88, 89], were mostly down-regulated after 24 hours. Overall, the differential regulation of genes in the gut involved in immunity, peptidase activity, and monooxygenase activity 24 hours after priming resembles the gene expression patterns found in whole larvae shortly after *Btt* exposure in other work [19, 31].

Five days after priming, guts showed additional stress in the ER and the degradation of ribosomes (Fig 7D). An even more notable phenomenon was the formation of enlarged microvilli (MV) heads (compare Fig 7B, E vs. 7C, F). Such MV “blebs” have been described in rats and can contain catalytically active enzymes with regulatory properties against microbes [90–92]. If and how MV “blebs” could affect *Btt* proliferation upon exposure to the spore diets is still unclear and an interesting question to follow up. Despite the obvious ultrastructural differences between the guts of primed and control larvae, the gene expression profiles were more similar compared to the early time points. Most DEGS were downregulated in primed guts (Fig 3A), including several genes encoding for proteins involved in the cytoskeletal structure of cells, which correlates with the aberrant appearance of the MV (Fig 7G). Also, several transmembrane transporter genes were down-regulated (Figs 7H and 4B). Similar transporters are reported to function as receptors for Cry toxins. However, the reported putative functional receptors for *T. castaneum* were unaffected [93–95]. Genes encoding for monooxygenases were mostly downregulated after 24 hours. As detoxifying enzymes are costly to maintain and, therefore, under strict regulation [96], the lower expression level in primed larval guts after 24 hours might have persisted to 5 days after.

The guts of primed and control larvae hardly differed on the ultrastructural level after exposure to infectious *Btt* spores (Fig 8A-F). In both cases, the gut epithelium was damaged, and vegetative cells of *Btt* were visible. The microvilli rim was degraded, and we found many apoptotic cells within the epithelia of primed and control guts. Overall, the effects of the infection seem so strong that they override smaller, priming-associated differences. Also, the number of DEGs was the lowest upon exposure to life *Btt* (Fig 3A). However, most of these DEGs were upregulated in primed guts (Fig 3A). Interestingly, many of their gene products have been described to bind to pathogen-derived molecules and invoke immune signaling. Examples are genes encoding for apolipophorinI/II, proteins containing leucine-rich repeats, odorant binding receptors, and a scavenger receptor [97–102]. Some of these genes have also been reported to be upregulated in *T. castaneum* upon *Btt* exposure in other studies [31, 103]. A gene encoding for Arylphorin, which has been correlated with *Bt* resistance in a Lepidopteran host by acting as a mitogen, was the second highest upregulated gene after *Btt* exposure in primed larvae compared to control larvae [104, 105]. In contrast to 24 hours after priming and 5 days after priming without exposure to *Btt* spores, monooxygenases, a Glutathion-S-transferase, and a Peroxiredoxin were upregulated in primed larvae after exposure to *Btt* spores. Overall, four hours after *Btt* spore exposure, primed larvae possessed an increased abundance of transcripts for genes involved in the binding and detoxification of foreign/pathogen compounds and cellular regeneration, which could imply a strategy to prevent damage done by the virulence factors of *Btt*.

For the first time, we described in detail the changes in the gut of *T. castaneum* larvae upon oral immune priming. In mosquitos and bean bugs, the microbiome breaches the epithelial cell lining, thereby influencing immune cell differentiation [15, 27]. Indeed, the microbiome in our system is needed for successful oral immune priming as well [26], and a change in its composition is described upon priming [28]. However, we here could not detect any signs of bacterial breaching in the gut epithelia on the ultrastructural level upon priming, and it is debated whether *Btt* breaches the gut before host death [106]. Pinaud *et al.* [17] showed that in snails, oral immune priming leads to a shift from a cellular immune response upon the first encounter with the parasite to a humoral response after the second encounter. In their system, however, the *Schistosoma* parasite forms sporocysts that develop within the host tissue, representing a different pathogenesis than that of *Btt*. These works highlight the differences in the investigated host–parasite models and reflect the diversity of immune priming phenomena in invertebrates [107]. In our case, virulence factors in the sterile filtered spent culture media supernatant of *Btt* induce a stress state in the gut of *T. castaneum* that resembles an actual infection at the ultrastructural (compare Figs 5A-F + 6A-F + 8A-F) and gene expression level. This stress state initially induces an immune response that diminishes over time, but upon re-infection with live *Btt* spores results in up-regulation of protective genes against *Btt*. Greenwood *et al.* [31] found an increased immune response in primed *T. castaneum* larvae 6 hours after exposure to life *Btt* using whole larvae. It is known that *T. castaneum* is able to build immune memory upon septic infection [14, 16]. Thus, investigating the potential crosstalk between the gut epithelium and the hemocoelic side would be interesting. The identified changes in the gut might induce hemocyte migration or differentiation or influence the gene expression profile of fat body tissue. This could lead to systemic immune responses and memory, which might aid protection against oral *Btt* infection and potentially also septic infection.

With respect to the five immune memory criteria as worked out by Pradeu and Du Pasquier [108], immune priming in *T. castaneum* meets the following: it is specific to a certain degree both via the septic and oral route [14, 18], it might enable a faster response as indicated by the early up-regulation of protective genes in the gut found in this study as well as the up-regulation of immune genes at whole larval level found by Greenwood et al. [31]. Protection in orally primed larvae lasts for at least 5 days, as shown in this study, and up to 8 days, as in some septic studies, and can be transmitted to subsequent generations [14, 23]. The initial priming response in the gut diminishes the gene expression level over five days, indicating extinction (Fig 3A + B).

But how could memory be formed in the insect gut epithelium? Work by Liu X et al. [81] in *Drosophila* showed that pathogenic microbes, in contrast to the normal microbiota, activate additional molecular pathways. This causes stem cells to differentiate into enteroendocrine cell types, changing the epithelial cell composition in the gut epithelium. Such changes might provide a type of memory that is based on epithelial cell structure. Accordingly, our WGCNA analysis identified differences in gene expression in the cell cycle and related processes 24 hours after priming (Fig 4A). Also, it is known from other systems that inflammatory memory exists in epithelial tissue in the form of chromatin rearrangements [5, 6]. Interesting candidates for the induction of memory and specificity of oral immune priming in our system are toxins such as Cry3Aa that are unique to certain strains of *Bt*. The damage done by these toxins, combined with the shifts in microbiome composition, could provide the stimuli needed to invoke epigenetic or architectural changes in the gut epithelium that lead to memory in non-immune tissue.

On this basis, the insect model might be important for future work on immune memory, which is not simply understood as a property of innate or adaptive immune cells in isolation. It seems rather a property of interacting cells and tissues that, on their own or in combination, might be able to form memory.

## Acknowledgements

We thank Kathrin Brüggemann, Ilka Rauch and Anke Große Brinkhaus for their technical assistance.

## Funding

We acknowledge funding by the Deutsche Forschungsgemeinschaft (DFG, German Research Foundation) within the Research Training Group GRK 2220 “Evolutionary Processes in Adaptation and Disease,” project number 281125614 to JK and partial funding within CRC TRR 212 (NC³) – project number 316099922 to JK.

## Supporting information captions

**S1 Fig: Survival of *T. castaneum* larvae exposed to either control or priming diets from the EM and RNA-seq experiments. A** Kaplan Meier curves displaying the survival of *T. castaneum* larvae exposed to either control diets (n =96 and 121), *Bt* medium diets (n =95 and 123) or priming diets (n =96 and 123) upon exposure to infectious *Btt* spores. **B** Forest plot representing the estimated fixed-effects coefficients of primed and control compared to medium (black, dashed line at 1.0).

**S2 Fig: Principal component analysis.** 3-dimensional PCA for differentially expressed genes identified with DESeq2. PC1 represents 40.34% variation, PC2 represents 15.25% variation and PC3 represents 9.66% variation.

**S3 Fig: Hierarchical cluster tree of the weighted gene co-expression analysis.** Each branch represents a single gene. Genes were clustered into modules based on their expression (unmerged), and closely related modules were merged based on their similarity in eigengene values (merged). The resulting modules are displayed in colors but were converted to numbers for later analysis.

**S4 Fig: Pearson correlation of module eigengene values with control or priming treatments at the different timepoints.** Cell color = pearson correlation coefficient. Asterisks = Significance of Student asymptotic p-value calculation (* if p < 0.05, ** if p < 0.01, *** if p < 0.001).

**S5 Fig: Volcano plots for proteomic analysis. A** Volcano plot for Btt vs Btt -cry supernatants without imputation for missing values. Plotted is the limma result for all protein groups that are at least in 3 of 4 samples of either Btt- or Btt -cry supernatants. X-axis represents the log2-fold-change (log2FC), the y-axis represents the negative log10-adjusted pvalue (−log10(adj.P.Val). The lines in the plot represent the chosen significance levels: The vertical lines are drawn at log2FC = −1/1, the horizontal line is drawn at −log10(0.05) = 1.301. Light green = significantly enriched protein groups. **. B** Volcano plot for Btt vs Btt -cry supernatants with imputation for missing values. Plotted is the limma result for th imputation of all protein groups that are at least in 3 of 4 samples of either Btt- or Btt -cry supernatants. X-axis represents the log2-fold-change (log2FC), the y-axis represents the negative log10-adjusted pvalue (−log10(adj.P.Val)). The lines in the plot represent the chosen significance levels: The vertical lines are drawn at log2FC = −1/1, the horizontal line is drawn at −log10(0.05) = 1.301. Light green = significantly enriched protein groups.

**S1 Table: Survival and larval growth measurement data.**

**S2 Table: Raw count data from RNA sequencing.**

**S3 Table: Meta information about Samples from the RNA-Sequencing run.**

**S4 Table: Result for the differential gene expression analysis using DESeq2.** Includes the results for the DESeq2 analyses at each timepoint between primed and control. Additionally contains categories used to group genes in biological context in the manuscript and the GO terms from the Clusterprofiler analyses.

**S5 Table: Exemplary proteomics results for supernatants derived from *Btt*** *Δ* ***p188* (C) and Btt (P) cultures. A** Log2 transformed label-free quantification (LFQ) values for candidate protein groups. **B** LFQ values upon imputation using the R package “imputeLCMD”. **C** Limma statistics results for imputed values. (Full table in S6 Dataset).

**S6 Table: Full proteomics table before and after imputation and Limma statistical analysis.**

**S1 Code: R Codes used for data analyses and figures.**

## Data availability

Raw data for the RNA-sequencing experiment can be found at: https://dataview.ncbi.nlm.nih.gov/object/PRJNA1174285?reviewer=1tvk9cnufn6k0qsbp3dqs2f56j. This data will be made publicly available under the Bioproject: PRJNA1174285 upon publication of the manuscript.

Raw data for the proteomics experiment can be found at https://repository.jpostdb.org/preview/68712976967221def02b9b. This data will be made publicly available under the identifier JPST003396 upon publication of the manuscript.

## Notes

### Competing Interest Statement

The authors have declared no competing interest.

### Summary of Updates

Reviewer access key for proteomics data removed. Supplementary material added.

