## Supplementary material (S1-S5 Figures, S5 Table) for "Immune priming in the insect gut: a dynamic response revealed by ultrastructural and transcriptomic changes"

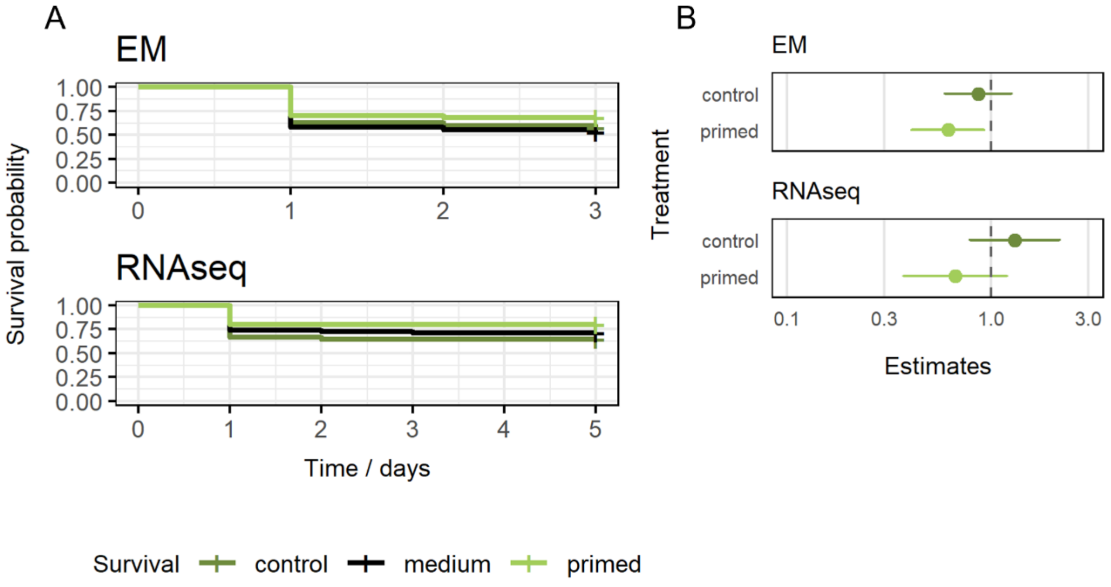


**S1 Fig: Survival of *T. castaneum* larvae exposed to either control or priming diets from the EM and RNA-seq experiments. A** Kaplan Meier curves displaying the survival of *T. castaneum* larvae exposed to either control diets (n =96 and 121), *Bt* medium diets (n =95 and 123) or priming diets (n =96 and 123) upon exposure to infectious *Btt* spores. **B** Forest plot representing the estimated fixed-effects coefficients of primed and control compared to medium (black, dashed line at 1.0).

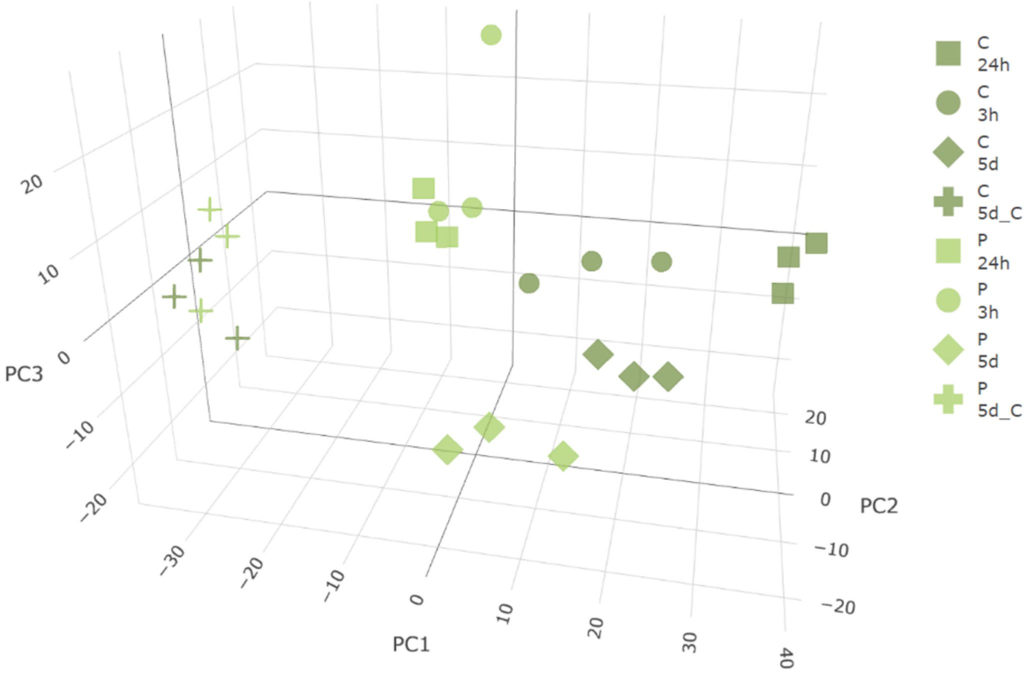


**S2 Fig: Principal component analysis.** 3-dimensional PCA for differentially expressed genes identified with DESeq2. PC1 represents 40.34% variation, PC2 represents 15.25% variation and PC3 represents 9.66% variation.

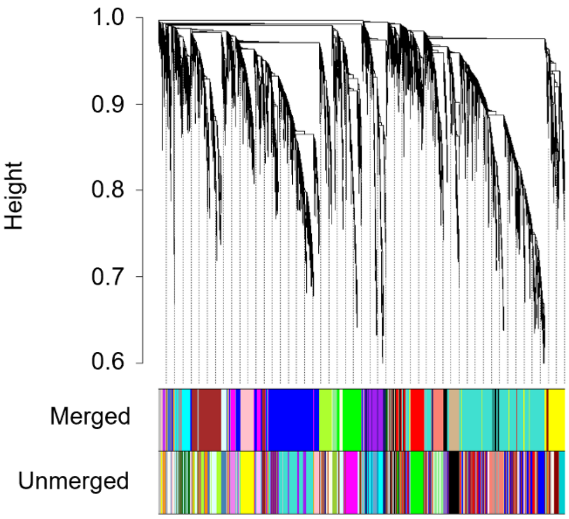


**S3 Fig: Hierarchical cluster tree of the weighted gene co-expression analysis.** Each branch represents a single gene. Genes were clustered into modules based on their expression (unmerged), and closely related modules were merged based on their similarity in eigengene values (merged). The resulting modules are displayed in colors but were converted to numbers for later analysis.

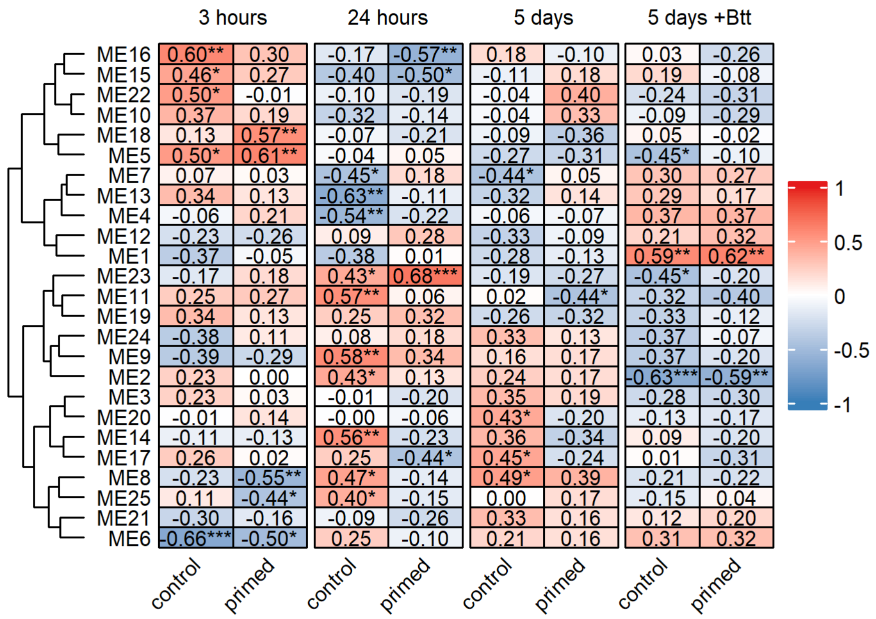


**S4 Fig:** Pearson correlation of module eigengene values with control or priming treatments at the different timepoints. Cell color = pearson correlation coefficient. Asterisks = Significance of Student asymptotic p-value calculation (* if p < 0.05, ** if p < 0.01, *** if p < 0.001).

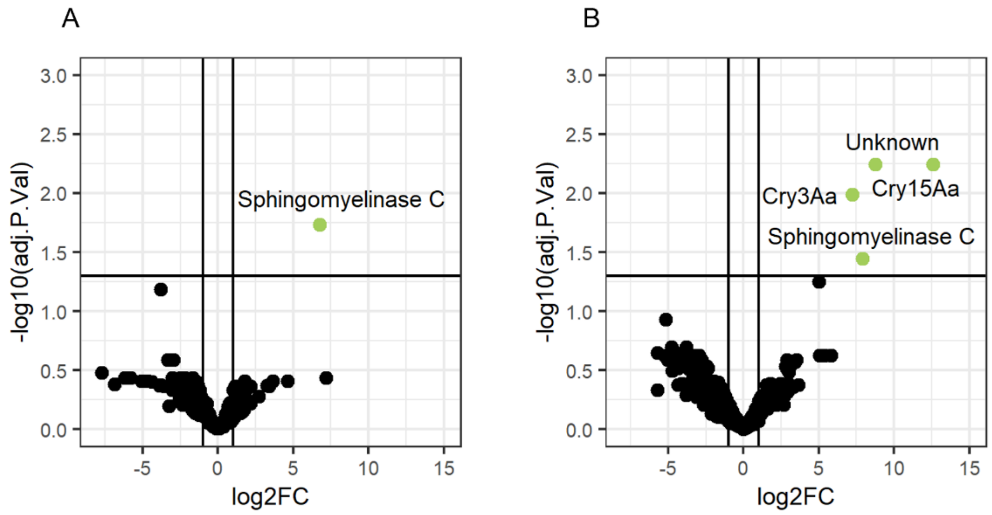


**S5 Fig: Volcano plots for proteomic analysis. A** Volcano plot for Btt vs Btt -cry supernatants without imputation for missing values. Plotted is the limma result for all protein groups that are at least in 3 of 4 samples of either Btt- or Btt -cry supernatants. X-axis represents the log2-fold-change (log2FC), the y-axis represents the negative log10-adjusted pvalue (-log10(adj.P.Val). The lines in the plot represent the chosen significance levels: The vertical lines are drawn at log2FC = -1/1, the horizontal line is drawn at -log10(0.05) = 1.301. Light green = significantly enriched protein groups. **. B** Volcano plot for Btt vs Btt -cry supernatants with imputation for missing values. Plotted is the limma result for th imputation of all protein groups that are at least in 3 of 4 samples of either Btt- or Btt -cry supernatants. X-axis represents the log2-fold-change (log2FC), the y-axis represents the negative log10-adjusted pvalue (-log10(adj.P.Val)). The lines in the plot represent the chosen significance levels: The vertical lines are drawn at log2FC = -1/1, the horizontal line is drawn at -log10(0.05) = 1.301. Light green = significantly enriched protein groups.

**S5 Table: Exemplary proteomics results for supernatants derived from *Btt $\Delta$p188* (C) and Btt (P) cultures. A** Log2 transformed label-free quantification (LFQ) values for candidate protein groups. **B** LFQ values upon imputation using the R package "imputeLCMD". **C** Limma statistics results for imputed values. (Full table in supplementary file ST2).

A

| Gene product | Log2(LFQ C-1) | Log2(LFQ C-2) | Log2(LFQ C-3) | Log2(LFQ C-4) | Log2(LFQ P-1) | Log2(LFQ P-2) | Log2(LFQ P-3) | Log2(LFQ P-4) |
|---|---|---|---|---|---|---|---|---|
| Cry15Aa | NA | NA | NA | NA | 31.01045322 | 31.44421028 | 33.40349776 | 30.67791493 |
| Unknown | NA | NA | NA | NA | 30.13919304 | 30.44825352 | 29.39599179 | 31.21222336 |
| Cry3Aa | NA | NA | NA | NA | 27.90443351 | 28.14812044 | 26.63547214 | 27.70371816 |
| Sphingomyelinase C | 23.95082569 | 24.76739161 | NA | 24.30243031 | 30.87547525 | 30.90741654 | 32.37572934 | 31.44992667 |
| Chitinase D | 30.73432725 | 30.84640214 | 27.10339587 | 31.22647645 | 30.99334477 | 30.60884776 | 31.19060561 | 30.18801317 |

B

| Gene product | Imputed C-1 | Imputed C-2 | Imputed C-3 | Imputed C-4 | Imputed P-1 | Imputed P-2 | Imputed P-3 | Imputed P-4 |
|---|---|---|---|---|---|---|---|---|
| Cry15Aa | 16.18201015 | 17.65483805 | 21.63324881 | 20.69851368 | 31.01045322 | 31.44421028 | 33.40349776 | 30.67791493 |
| Unknown | 21.81732893 | 22.19854349 | 19.96800715 | 22.1940742 | 30.13919304 | 30.44825352 | 29.39599179 | 31.21222336 |
| Cry3Aa | 20.70763363 | 20.79312547 | 19.24434662 | 20.81952254 | 27.90443351 | 28.14812044 | 26.63547214 | 27.70371816 |
| Sphingomyelinase C | 23.95082569 | 24.76739161 | 20.91166129 | 24.30243031 | 30.87547525 | 30.90741654 | 32.37572934 | 31.44992667 |
| Chitinase D | 30.73432725 | 30.84640214 | 27.10339587 | 31.22647645 | 30.99334477 | 30.60884776 | 31.19060561 | 30.18801317 |

C

| Gene product | logFC | AveExpr | t | P.Value | adj.P.Val | B |
|---|---|---|---|---|---|---|
| Cry15Aa | 12.59186637 | 12.59186637 | 11.48649986 | 2.08856E-05 | 0.005671976 | 1.915561468 |
| Unknown | 8.754426984 | 8.754426984 | 10.6300036 | 3.30727E-05 | 0.005671976 | 1.711836963 |
| Cry3Aa | 7.206778999 | 7.206778999 | 8.968190846 | 8.95747E-05 | 0.010241373 | 1.206080418 |
| Sphingomyelinase C | 7.919059724 | 7.919059724 | 6.845816829 | 0.000416461 | 0.035711489 | 0.252255808 |
| Chitinase D | 0.767552399 | 0.767552399 | 0.681932528 | 0.519950977 | 0.768720626 | -5.670291331 |
